# Swift Large-scale Examination of Directed Genome Editing (SLEDGE Hammer)

**DOI:** 10.1101/479261

**Authors:** Omar T. Hammouda, Thomas Thumberger, Joachim Wittbrodt

**Affiliations:** Centre for Organismal Studies Heidelberg, Heidelberg University, Im Neuenheimer Feld 230, 69120 Heidelberg, Germany.; Heidelberg Biosciences International Graduate School, Heidelberg University, Im Neuenheimer Feld 501, 69120 Heidelberg, Germany.

## Abstract

In the era of CRISPR gene editing and genetic screening, there is an increasing demand for quick and reliable nucleic acid extraction pipelines for rapid genotyping of large and diverse sample sets. Despite continuous improvements of current workflows, the handling-time and material costs per sample remain the major limiting factors. Here we present a robust method for low-cost DIY-pipet tips addressing these needs; i.e. using a cellulose filter disc inserted into a regular pipet tip. These filter-in-tips allow for a rapid, stand-alone three-step genotyping workflow by simply binding the DNA contained in the primary lysate to the cellulose filter, washing it in water and eluting it directly into the buffer for the downstream application (e.g. PCR). This drastically cuts down processing time to maximum 30 seconds per sample, with the potential for parallelizing and automation. We show the ease and sensitivity of our procedure by genotyping genetically modified medaka and zebrafish embryos (targeted CRISPR/Cas9 knock-out and knock-in) in a 96-well plate format. The robust isolation and detection of multiple alleles of various abundancies in a mosaic genetic background allows phenotype-genotype correlation already in the injected generation, demonstrating the reliability and sensitivity of the filter-in-tips. Furthermore, our method is applicable across kingdoms with samples ranging from cells to tissues (e.g. plant seedlings, adult flies, mouse cell culture and tissue as well as adult fish fin-clips).

## Introduction

Gene-editing tools such as CRISPR/Cas9 have revolutionized genome editing in most model organisms (Cong et al., 2013; Mali et al., 2013; Stemmer et al., 2015). The easy application of this genome targeting method in organisms as well as cell culture allows extensive high-throughput applications such as genetic screens (Baranski et al., 2018; Kweon and Kim, 2018). Common to all genome targeting applications is the need for subsequent genotyping. Following PCR amplification of the targeted loci, among the most commonly used assays for indel detection are mismatch cleavage assays (using T7 Endonuclease I or Cel-based Surveyor assay), high-resolution melting analysis (HRMA) and sequencing (Thomsen et al., 2012; Zischewski et al., 2017). However, the high-throughput approaches face a bottle neck when it comes to genomic DNA extraction which usually requires a time and material consuming protocol.

Genome extraction has greatly improved due to the numerous advances on nucleic acid purification methods. These either rely on solid-phase extraction kits or on traditional phenol-chloroform purification, both of which are time or material consuming and require a lot of steps (Ali et al., 2017). Recent publications have shown the ability of membranes of different materials to bind nucleic acids with high affinity, thus eliminating the need for nucleic acid purification by direct amplification of the nucleic acid off the membrane (Jangam et al., 2009; Kim et al., 2010; McFall et al., 2015). A recent improvement has greatly reduced the genome extraction step to under 30 seconds using cellulose paper-based dipsticks (Zou et al., 2017). Although extremely advantageous in terms of time and material, these methods require fabrication or manufacturing steps, which make them unsuitable for high-throughput applications, especially for automated systems.

Teleost fish such as the Japanese rice fish medaka (*Oryzias latipes*) have been vertebrate model organisms for more than a century (Wittbrodt et al., 2002). Numerous advantages such as high fecundity and fast generation times; transparency of chorion and body allow for live non-invasive *in vivo* imaging; and the availability of various genetic tools makes them an attractive model organism for developmental, genetic and molecular studies (Kirchmaier et al., 2015). Moreover, the high efficiency of genome-editing in medaka by CRISPR/Cas9 renders it very attractive for high-throughput *in vivo* screening applications (Stemmer et al., 2015). However, despite the continuous development and progression of high-throughput automated imaging and analysis pipelines, current genotyping protocols display the limiting step.

In this study, we have developed a very fast, simple and efficient genotyping protocol, which has great potential for high-throughput and automated applications whenever it comes to nucleic acid extraction and transfer. In a medaka and zebrafish context, we describe our Swift Large-scale Examination of Directed Genome Editing (SLEDGE Hammer) protocol. Here we demonstrate rapid genotyping in a 96-well based format of various CRISPR/Cas9 edited medaka and zebrafish embryos. In brief, a cellulose paper disc inserted into conventional pipet tips allows for skipping purification of genomic DNA from cell/tissue lysates and facilitates direct use with PCR buffers. Our filter-in-tip approach allows easy high-throughput upscaling by using multichannel pipets to handle several samples simultaneously; while also offering potential for automated pipetting systems suitable for high-throughput automated screens. The filter-in-tips are generally applicable when transfer of nucleic acids is aimed for as shown by genomic DNA extraction from varying sources of tissues derived from diverse (model) organisms such as *Arabidopsis thaliana*, *Chironomus riparius*, *Drosophila melanogaster*, Mouse ear punches, Mouse Embryonic stem cells, fish embryos and fin-clips.

## Results and Discussion

Teleost fish have been extensively used for high-throughput approaches (Kithcart and MacRae, 2017; Lessman, 2011; Oxendine et al., 2006; Spomer et al., 2012) due to their small sized embryos, transparency and biological relevance as vertebrates. Recently, there have been various key developments in medaka research: 1) the improved efficiency of medaka genome editing via CRISPR/Cas9 (Stemmer et al., 2015); and 2) the establishment of the first wild vertebrate medaka inbred panel (Spivakov et al., 2014), which is the basis for phenotype / population genomic studies that rely on automated phenotyping as well as reliable and precise genotyping.

These developments allow us to foresee a surge in the use of medaka for various high-throughput assays (e.g. genetic screens, drug screens and Genome Wide Association Studies). However, genotyping a large number of specimens still remains an obstacle in terms of time and effort, as standard genotyping protocols start with grinding of embryos/tissue in lysis buffer and incubating for an extended period of time, followed by genomic DNA extraction and precipitation (most commonly using Isopropanol). Precipitation however comes with the risk of losing the genomic DNA sample. Taken together, this purification procedure can easily take 2 days and is hence not feasible in a high-throughput screening context.

In the light of the reported nucleic acid binding property of cellulose based filter paper (Zou et al., 2017), we sought to exploit this feature in order to by-pass the tedious genomic DNA purification step, thus saving time and material. To this end, we used a paper puncher to create Whatman filter paper discs (≈2 mm) and placed them inside of standard yellow 200 µl tips (filter-in-tips; Fig. 1A). To test the applicability of these modified pipet tips for rapid genotyping, the fins of 3 male and 3 female adults were clipped and administered to rapid genomic DNA extraction for sex determination by PCR. Fin clips were put into 100 µl fin-clip buffer-containing Eppendorf tubes. 50 µl of PCR mix with primers for the *non-muscle actin b* (*actb*) serving as control and *dnmtY* for genomic male sex determination were prepared for each specimen. After short grinding of the tissue, the filter-in-tips were used to soak the Whatman paper disc with the fin-clip lysate for the nucleic acids to bind to the cellulose simply by pipetting the liquid up and letting it rest in the tip for ≈10 seconds. Subsequently the lysate was released back into the tube for storage. Next, the filter was washed to remove lysate buffer reagents (which impair later diagnostics) by brief 1-2 washing steps in MilliQ water (individual aliquots per tip) by pipetting up and down. Finally soaking the filter disc for ≈5-10 seconds in the PCR reaction mix by pipetting up with the very same filter-in-tip released enough material for amplification. Whereas the *dnmtY* band correlated with male sex phenotype, *actb* control amplicon was present in all specimens (Fig. 1B).

**Figure 1.**
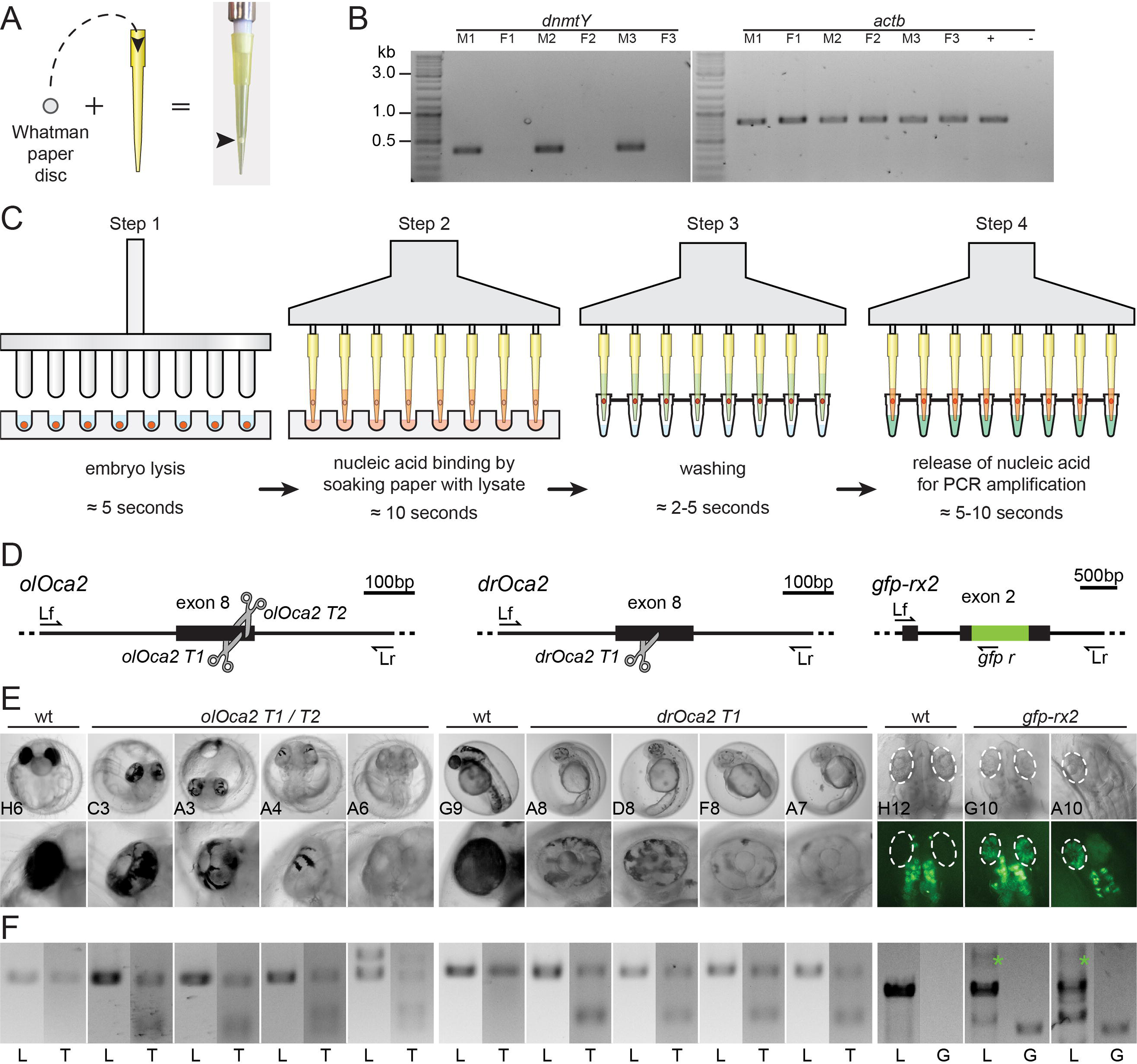
Filter-in-tips are highly sensitive for rapid transfer of nucleic acids for targeted genome editing diagnostics. A) Assembly of filter-in-tip – Whatman filter paper / cellulose disc inserted into 200 µl yellow tip at the sample-proximal end. B) Genomic male sex determination by *dnmtY* PCR of 3 male (M1-3) and 3 female (F1-3) fin clips. *actb* amplification as control, purified and precipitated wt DNA as positive control (+), water as negative control (-). Note: transfer of gDNA via filter-in-tips and specific amplification of loci was successful in all cases. C) Schematic workflow of high-throughput 96-well plate based SLEDGE-Hammer protocol with filter-in-tips. Step 1 Lysis: (96-pin) mortar used to simultaneously grind individual embryos in fin-clip buffer. Step 2 Binding: pipet up lysate to let nucleic acids (red) bind to cellulose filter discs (soak for ≈10 sec) using a multichannel pipet equipped with filter-in-tips. Release lysate back in wells for storage. Step 3 Washing: wash filter discs containing nucleic acids (red) by pipetting nuclease free water in (wait ≈2-5 sec) and out. Step 4 Elution: pipet up pre-mixed PCR mixture (wait ≈5-10 sec) to release nucleic acids and pipet back for amplification. D) Schematic representation of CRISPR/Cas9 mediated NHEJ-based knock-out (medaka *oca2*, *olOca2*; zebrafish *oca2*, *drOca2*) and HDR-mediated single-copy integration of *gfp* sequence in frame with medaka *rx2* (cf. Gutierrez et al., 2018 (Gutierrez-Triana et al., 2018)). Location of sgRNA target sites shown by scissors. Primers used for analysis indicated; locus forward (Lf), locus reverse (Lr) of the respective gene. *rx2* locus shown after single-copy HDR-mediated integration of *gfp*. E) Besides fully pigmented eyes in wt medaka and zebrafish, varying degrees of pigment loss in body and RPE (blow-up) of representative crispants injected with individual sgRNAs targeting *oca2* exon 8. Retinae (dashed ellipses) of injected embryos show GFP expressing cells after HDR/donor mediated integration of *gfp* into *rx2* open reading frame. Note: unspecific autofluorescence of body pigment. 96-well plate coordinates of specimens indicated (see Supplementary Fig. S1). F) Genotyping of representative embryos in E. Locus PCR (L; primers Lf/Lr) of respective genes and T7EI assay (T) validating indel formation in *oca2* loci. In *gfp-rx2* tagging, besides non-*gfp*-integrated locus band (L; primers Lf/Lr), single *gfp* integration evident by PCR (green asterisk). Integration validated by locus-gfp band (G; primers Lf/*gfp* r).

We next raised the question if this SLEDGE-Hammer protocol is robust and reliable for large scale individual extractions of genomic DNA by equipping a multichannel pipettor with the filter-in-tips (Fig. 1 C). And further, if our protocol would allow to detect minor nucleotide changes as to be found in mosaic embryos resulting from CRISPR/Cas9-mediated knock-out. We thus targeted the *oculocutaneous albinism 2* (*oca2*, Fig. 1D) gene responsible for melanin pigmentation using three individual sgRNAs in combination with *Cas9* mRNA and *gfp* mRNA as injection marker in medaka and zebrafish one cell stages. Loss of pigmentation requires a bi-allelic mutation to occur in the *oca2* open reading frame, rendering this gene a perfect candidate for CRISPR/Cas9 knock-out efficiency analysis (Lischik et al., 2018). In addition to the knock-out attempts caused as a result of the error-prone non-homologous end-joining (NHEJ) repair mechanism, a more delicate CRISPR/Cas9 approach for tagging the *Retinal homeobox protein 2* (*Rx2*) gene with *gfp* was used (Fig. 1D). A biotinylated dsDNA PCR donor fragment comprising the *gfp* sequence flanked by nucleotide stretches homologous to about 400bp up and downstream of the *rx2* sgRNA target site was co-injected with *Cas9* mRNA and the respective *rx2* sgRNA (Gutierrez-Triana et al., 2018). Per *oca2* sgRNA, 21 positively injected embryos (4 days post fertilization (dpf) for medaka, 2 dpf for zebrafish) were randomly selected and put into a 96 U-well plate for phenotype/genotype screening. For screening of *gfp-rx2* integration, 21 injected embryos were randomly picked at 2 dpf and administered alike (Supplementary Fig. S1A). One negative control (empty well containing embryo rearing medium) as well as two uninjected controls complemented each group.

Phenotyping revealed that the knock-out attempts in the *oca2* genes caused the expected loss of pigmentation in all specimens with various extents as evident in the retinal pigmented epithelium (RPE; Fig. 1E). Tagging of the *rx2* gene was visible by specific GFP expression in the medaka retinae (Fig. 1E). HDR-mediated knock-in attempts are less efficient than knock-out approaches, still ≈40 % (8/21) of injected specimens showed varying numbers of GFP positive cells in the developing eye. To correlate with the observed phenotype, each embryo was genotyped individually.

Based on the 96 well plate format used for long term single-embryo monitoring / phenotyping, we additionally have created a stainless-steel 96-pin mortar (a.k.a. ‘the Hammer’). After phenotyping (plate imaging), the fish medium in the 96 U-well plate was replaced with fin-clip lysis buffer (100 µl per well) and the embryos were simultaneously ground using our custom-made mortar. For individual genotyping we followed the rapid protocol described above using the filter-in-tips in combination with a multichannel pipet (Fig. 1C). For the *oca2* knock-out embryos, locus amplification revealed that in all cases transfer of genomic DNA to the PCR mixture was successful (Fig. 1F, Supplementary Fig. S1B). We subsequently examined the success of targeted genome editing (indel formation) in the *oca2* loci using T7 Endonuclease I (T7EI) assay, which detects and cuts heteroduplexes caused by a heterogeneous mixture of indel alleles from genome targeted specimens. T7EI genotyping confirmed the phenotypic observation, i.e. all injected embryos of all *oca2* knock-out attempts showed heteroduplex digestion in the T7EI assay, whereas the locus amplicon of the respective uninjected controls did not (Fig. 1F, Supplementary Fig. S1C). Furthermore, all *gfp-rx2* tagging approaches that were positive in the GFP screening could as well be confirmed by PCR genotyping as fusion of the *gfp* sequence with the *rx2* locus in the respective specimens (Fig. 1F, Supplementary Fig. S1D-E). It is interesting to note that in 7 out of the 8 positively screened embryos, a PCR band indicative for single-copy HDR-mediated integration was evident (Fig. 1F, Supplementary Fig. S1D) whereas the integration of the donor cassette probably underwent NHEJ-mediated concatenation and integration in four cases (Supplementary Fig. S1E).

Besides genotype/phenotype confirmation, all PCRs and T7EI assay revealed that the filter-in-tip approach robustly and reliably extracted genomic DNA for PCR amplification and that the transferred genomic DNA contained enough allele variants for the subsequent T7EI assay. Furthermore, considering genotyping of 96 embryos, conventional genomic DNA extraction could easily have taken 2 days and would require tedious and repetitive pipetting. In contrast, our SLEDGE-Hammer protocol allows drastic shortening of time and pipetting steps to a maximum of 10-15 minutes per plate using a standard 8-channel multipipettor. This effortless genotyping will massively upscale with the continuous advancements of automated multipipettors, which have now reached up to 384 channels.

Besides the reliable extraction of genomic DNA from fish embryos, we investigated if this method can be generalized for genomic DNA extraction from varying sources of tissues derived from (model) organisms such as *Arabidopsis thaliana*, *Drosophila melanogaster*, *Chironomous riparius*, mouse ear punches (EP) and mouse embryonic stem cells (mESC) (Fig. 2). In all cases, grinding of tissue, i.e. a single seedling of Arabidopsis, an individual adult fly of both insect species as well as an individual mouse ear punch and 2.5 × 10^5^ ESC suspension in lysis buffer and subsequent transfer of genomic DNA by pipetting of the lysate using our filter-in-tips transferred enough material for specific locus amplification.

**Figure 2.**
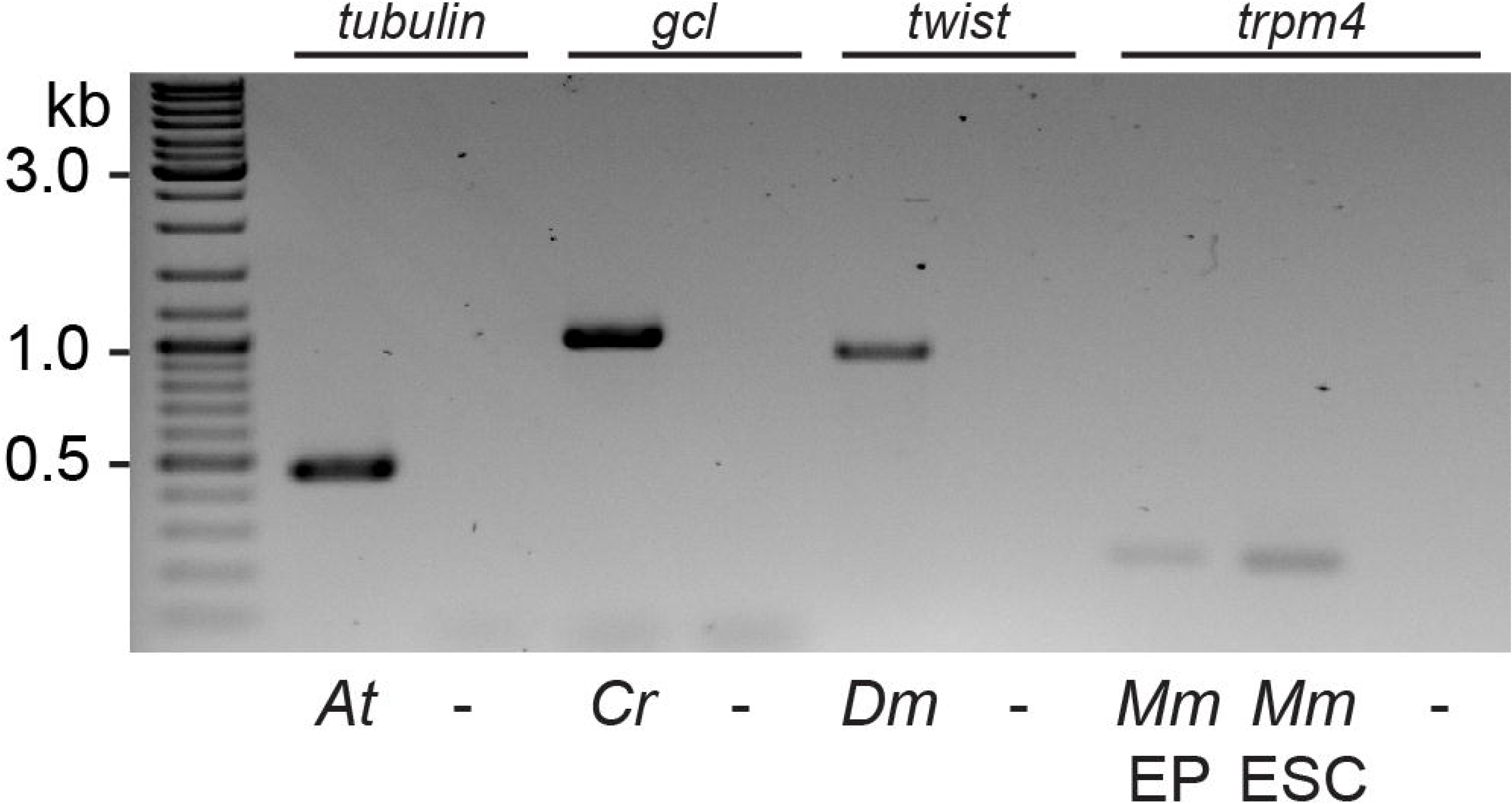
General applicability of filter-in-tips to rapidly transfer genomic DNA from various sources. Filter-in-tips successfully transferred genomic DNA from lysates of *Arabidopsis thaliana* (*At*), *Chironomus riparius* (*Cr*), *Drosophila melanogaster* (*Dm*), mouse (*Mus musculus*, *Mm*) ear punches (EP) and mouse Embryonic Stem Cells (ESCs) as evident by PCR amplification of candidate genes (*tubulin*; *germ cell-less*, *gcl*; *twist*; *trpm4*). Water control (-).

Taken together, our SLEDGE Hammer protocol with the adapted filter-in pipet tips, allows to bypass the otherwise tedious and time-consuming genomic purification step that hitherto limited high-throughput genotyping approaches. With the rapid genotyping method presented here, future attempts of individual phenotype-genotype correlation following high-throughput automated screening of individual specimens is in reach. Furthermore, our simple modification of already available pipet tips can easily be adapted for use in fully automated research applications involving almost any biological sample. Due to the drastic reduction in time and costs, we further foresee this application for stock maintenance in laboratories or stock centers, where genotyping is a daily routine.

## Materials and Methods

### Ethics Statement

All fish are maintained in closed stocks at Heidelberg University. Medaka (*Oryzias latipes*) and zebrafish (*Danio rerio*) husbandry (permit number 35–9185.64/BH Wittbrodt) were performed according to local animal welfare standards (Tierschutzgesetz §11, Abs. 1, Nr. 1) in accordance with European Union animal welfare guidelines (Bert et al., 2016). The fish facility is under the supervision of the local representative of the animal welfare agency. Embryos of medaka of the wildtype Cab strain and of zebrafish AB/AB line were used at stages prior to hatching. Medaka were raised and maintained as described previously (Köster et al., 1997).

### sgRNA target site selection and *in vitro* transcription

sgRNAs were designed with CCTop as described in Stemmer et al. (Stemmer et al., 2015). sgRNA for *rx2* was the same as in Stemmer et al. (Stemmer et al., 2015) and the *olOca2* sgRNAs were the same as in Lischik et al. (Lischik et al., 2018). The following target sites were used (PAM in brackets): *rx2 T1* (GCATTTGTCAATGGATACCC[TGG]), *olOca2 T1* (GAAACCCAGGTGGCCATTGC[AGG]), *olOca2 T2* (TTGCAGGAATCATTCTGTGT[GGG]), *drOca2 T1* (GTACAGCGACTGGTTAGTCA[TGG]). Cloning of sgRNA templates and *in vitro* transcription was performed as detailed in Stemmer, et al. (Stemmer et al., 2015).

### Microinjection

Medaka one-cell stage embryos were injected in the cytoplasm as previously described (Stemmer et al., 2015), zebrafish one-cell stage embryos were injected in the cytoplasm or yolk. Injection solutions for *oca2* targeting comprised: 150 ng/µl *Cas9* mRNA, 15 ng/µl respective sgRNA (either *olOca2 T1, olOca2 T2* or *drOca2 T1*) and 10 ng/µl *gfp* mRNA as injection tracer. Injected embryos were incubated at 28 °C and screened for GFP expression at 1 dpf. Injection solution for *gfp-rx2* tagging comprised 150 ng/µl Cas9 mRNA, 15 ng/µl *rx2 T1* sgRNA and 5 ng/µl biotinylated PCR donor fragment (Gutierrez-Triana et al., 2018).

### Sample preparation and Imaging

Medaka and zebrafish embryos were administered to a 96 U-well microtiter plate (Nunc, Thermofisher #268152) containing either 1x embryo rearing medium or 1x zebrafish medium for automated screening/phenotyping using an ACQUIFER Imaging Machine (DITABIS AG, Pforzheim, Germany). Images were acquired in brightfield using 9 z-slices (dz = 100 µm) and a 4x Plan UW N.A. 0.06 (Nikon, Düsseldorf, Germany) to capture the centered embryo. Integration times were fixed with 100 % relative white LED intensity and 50 ms exposure time. GFP channel was used on *rx2* specimens at 30 % relative LED intensity and 200 ms exposure time.

### Nucleic acid extraction and locus amplification by PCR

Fin-Clip Lysis Buffer was used for medaka, zebrafish, *A. thaliana*, *C. riparius* and *D. melanogaster* (0.4 M Tris-HCl pH 8.0, 5 mM EDTA pH 8.0, 0.15 M NaCl, 0.1 % SDS in Milli-Q water). For *M. musculus* ear clip and embryonic stem cell samples 1 % SDS Lysis Buffer was used (0.1 M Tris-HCl pH 8.0, 5 mM EDTA pH 8.0, 0.2 M NaCl, 1 % SDS in Milli-Q water).

For medaka and zebrafish extraction of genomic DNA, see main text. Our 96-well U-plate adapted stainless-steel 96-pin mortar was pre-cleaned by incubation in hypochlorite solution (1:10 dilution of commercial bleach reagent) for at least 15 minutes followed by 5 minutes incubation in MilliQ water. Plate with embryo lysate can be optionally stored at 4 °C with proper plate seal. 2 mm diameter paper puncher (Harris Uni-Core 2.0) was used on Whatman Cellulose paper (3030-917 Grade 3MM) to produce the paper discs which were transferred directly from the puncher to standard yellow 200 µl pipet tips (Steinbrenner GmbH). 50 µl PCR reaction mixes for the amplification of the desired genomic loci were prepared on ice using 1x Q5 reaction buffer, 200 µM dNTPs, 200 µM primer forward and reverse and 0.6 U/µl Q5 polymerase (New England Biolabs). 30 PCR cycles were run in all samples except for *A. thaliana*, *C. riparius* and *D. melanogaster* (35 cycles); annealing temperatures, extension times and primer sequences are given and PCR conditions used are listed in Table 1. Our filter-in-tips were used to transfer nucleic acids from tissue lysate to the PCR mix in three simple steps. 1) Binding: pipetting enough tissue lysate (here 50 µl) to soak the paper disc and wait ≈10 seconds before releasing the lysate back. 2) Washing: 1-2 washing steps by brief up and down pipetting in MilliQ water, the higher the concentration of SDS the more washing steps are required, i.e. for 1 % or higher SDS containing buffers, 4 washing steps are advised. 3) Elution: pipetting up the PCR mix to soak the paper disc (here 50 µl), wait ≈5-10 seconds and release the PCR mix which will contain enough nucleic acids to be amplified. Following PCR, 10 µl PCR reaction + 2 µl 6x orange Loading Dye were loaded on 1.5 % Agarose in 1xTAE gels. Gel electrophoresis was performed at 90V.

**Table 1.**
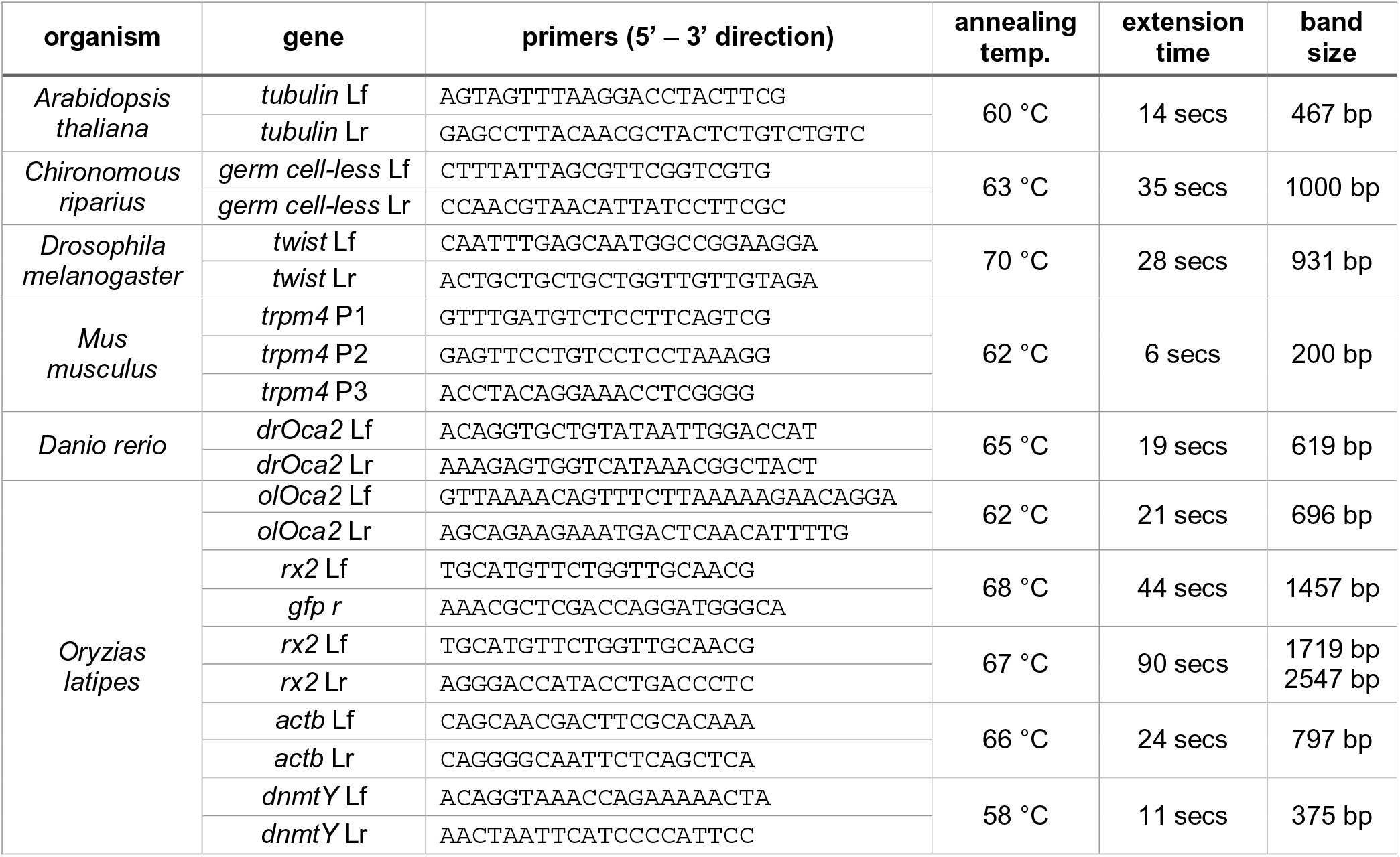
List of primers and PCR conditions used All primers used in this study are given in 5’-3’ direction. Expected band size displayed. Note: for *gfp-rx2* two major bands are expected upon HDR-mediated integration – the wt size of non-*gfp*-integrated loci as well as the larger single-*gfp* insertion. *Rx2* tagging design is as described in Gutierrez et al., 2018.

### Genotyping

Genotypes of samples used in our study were assessed based either on PCR band size or using the T7 Endonuclease I Assay (New England Biolabs). However, in order to save time, instead of purifying the PCR reactions, we directly transferred 10 µl of the unpurified PCR reaction into fresh PCR tubes with 7.5 µl Milli-Q water and 2 µl NEB Buffer 2 (total 19.5 µl) and proceeded with heteroduplex formation (1: Denature at 95 °C for 5 mins, 2: Stepwise cool down from 95 °C to 85 °C with −2 °C/sec, and 3: Stepwise cool down from 85 °C to 25 °C with −0.1 °C/sec) and subsequent digestion by adding 0.5 µl of T7EI (NEB) and incubation for 30 min at 37 °C. After incubation, the digest was immediately run on a 1.5 % Agarose in 1xTAE gel at 90 V.

## Supporting information

## Acknowledgments

We are grateful to F. Böttger for building a working copy of ‘Thor’s Hammer’ for this genotyping approach. Further we like to thank S. Lemke for Drosophila and Chironomous flies and primers, M. Freichel for the mouse ear punch and primers, L. Zilova for the mouse ES cells and J. Lohmann for the Arabidopsis seedlings and primers. We further thank all members of the Wittbrodt lab for their critical, constructive feedback on the procedure and the manuscript. OH, member of the Heidelberg International Biosciences Graduate School (HBIGS), was supported by a fellowship of the DZHK. This work was supported by the coordinated program of the DFG (FOR2509, project P10).

## Author contribution

O.H., T.T. and J.W. designed the study, analysed the data and wrote the manuscript. O.H. and T.T. performed injection experiments and acquired the data. O.H. performed all genotyping and indel assay analyses.

**Supplementary Figure 1. 96-well format high-throughput SLEDGE-Hammer analysis**

A) 96 well plate layout for high-throughput genotyping. CRISPR/Cas9 mediated (green wells) knock-out of *oca2* locus with three individual sgRNAs: *olOca2 T1* (columns 1-3), *olOca2* T2 (columns 4-6), *drOca2 T1* (columns 7-9). HDR/donor mediated integration of *gfp* in frame with *rx2* locus (columns 10-12). Medium control (blue wells) and uninjected wildtype specimens (red wells) included for control. B) Successful rapid extraction/transfer of genomic DNA using filter-in-tips evident by *oca2* locus PCR amplification of injected and uninjected specimens. Larger random indel formation can yield extra bands (black asterisks). C) T7EI assay of locus amplification in B reveals specificity of genomic DNA transfer method by T7EI digestion of heteroduplexes (cut bands) in *oca2* crispants but not wildtype embryos. D) *rx2* locus PCR amplification of injected and uninjected specimens. Note: non-*gfp*-integrated locus band (black arrowhead, 1719 bp) and single precise *gfp* integration (green asterisks, 2547 bp) evident by band-size. Additional bands stem from NHEJ-events. E) *gfp-rx2* specific bands correlate with embryos expressing GFP in retinae. All single-copy HDR-mediated *gfp* integration events in D could as well be verified by band size (953 bp) here. In addition, some donors underwent NHEJ (red asterisk, ≈1400 bp) or most probable concatenation events (yellow asterisks).

