## Supplementary material for "Swift Large-scale Examination of Directed Genome Editing (SLEDGE Hammer)"

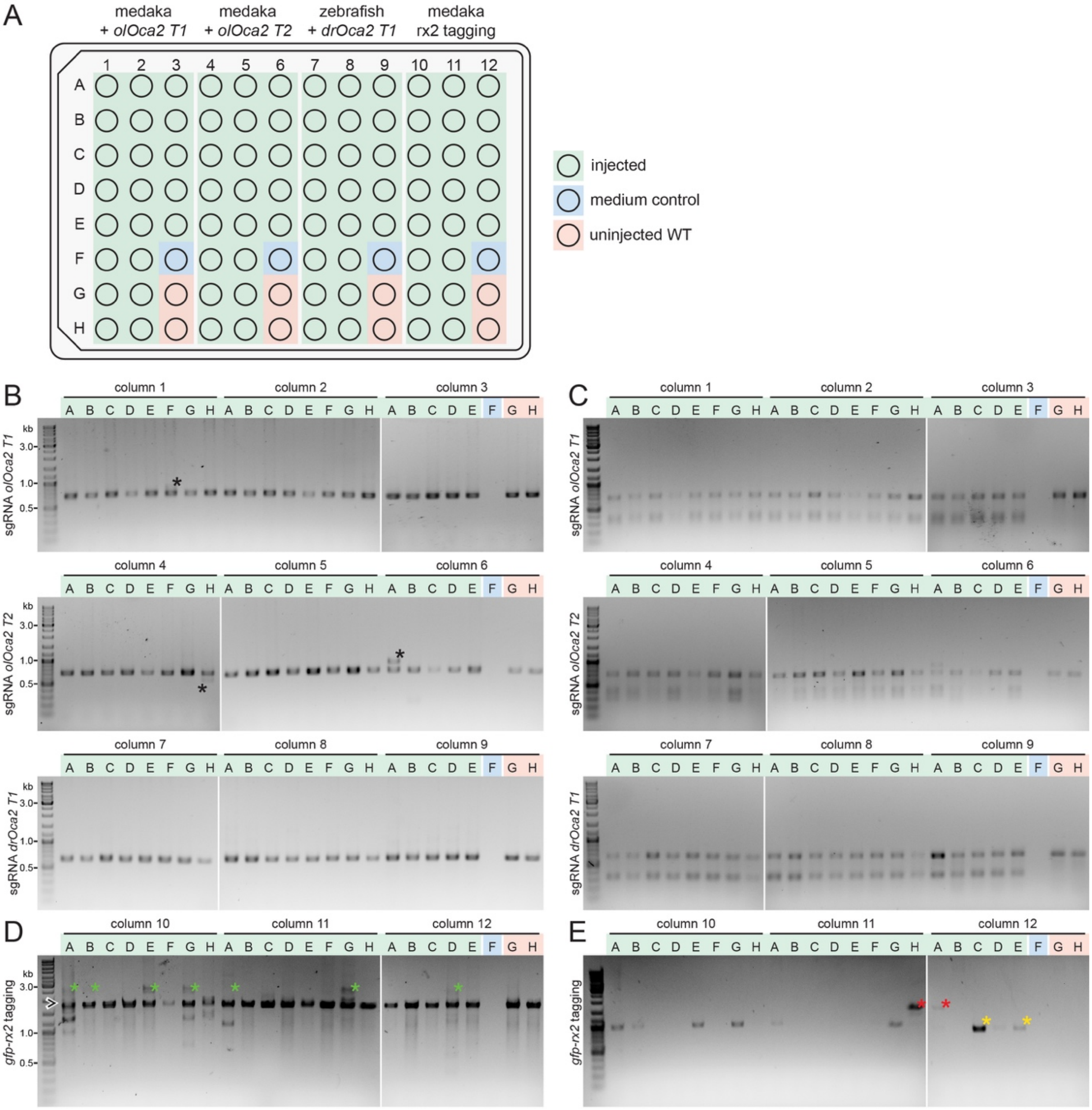
A
medaka + olOca2 T1
medaka + olOca2 T2
zebrafish + drOca2 T1
medaka rx2 tagging
injected
medium control
uninjected WT
B
C
D
E
column 1
column 2
column 3
column 4
column 5
column 6
column 7
column 8
column 9
column 10
column 11
column 12
sgRNA olOca2 T1
sgRNA olOca2 T2
sgRNA drOca2 T1
gfp-rx2 tagging
kb
3.0
1.0
0.5

**Supplementary Figure 1. 96-well format high-throughput SLEDGE-Hammer analysis**

A) 96 well plate layout for high-throughput genotyping. CRISPR/Cas9 mediated (green wells) knock-out of *oca2* locus with three individual sgRNAs: *olOca2 T1* (columns 1-3), *olOca2* T2 (columns 4-6), *drOca2 T1* (columns 7-9). HDR/donor mediated integration of *gfp* in frame with *rx2* locus (columns 10-12). Medium control (blue wells) and uninjected wildtype specimens (red wells) included for control. B) Successful rapid extraction/transfer of genomic DNA using filter-in-tips evident by *oca2* locus PCR amplification of injected and uninjected specimens. Larger random indel formation can yield extra bands (black asterisks). C) T7EI assay of locus amplification in B reveals specificity of genomic DNA transfer method by T7EI digestion of heteroduplexes (cut bands) in *oca2* crispants but not wildtype embryos. D) *rx2* locus PCR amplification of injected and uninjected specimens. Note: non-*gfp*-integrated locus band (black arrowhead, 1719 bp) and single precise *gfp* integration (green asterisks, 2547 bp) evident by band-size. Additional bands stem from NHEJ-events. E) *gfp*-*rx2* specific bands correlate with embryos expressing GFP in retinae. All single-copy HDR-mediated *gfp* integration events in D could as well be verified by band size (953 bp) here. In addition, some donors underwent NHEJ (red asterisk, ≈1400 bp) or most probable concatenation events (yellow asterisks).
